# Structural organization of the C1b projection within the ciliary central apparatus

**DOI:** 10.1101/2021.06.16.448709

**Authors:** Kai Cai, Yanhe Zhao, Lei Zhao, Nhan Phan, George B. Witman, Daniela Nicastro

**Affiliations:** Departments of Cell Biology and Biophysics, University of Texas Southwestern Medical Center, Dallas, Texas 75231, USA; Division of Cell Biology and Imaging, Department of Radiology, University of Massachusetts Medical School, Worcester, MA 01655, USA

**Author notes:** authors contributed equally (Yanhe Zhao, Lei Zhao). Institute of Crop Sciences, Chinese Academy of Agricultural Sciences, Beijing, 100081, PR China. Corresponding Author: Daniela Nicastro, Departments of Cell Biology and Biophysics, UT Southwestern Medical Center, Dallas, TX 75390, USA.

**Keywords:** Axoneme, ciliary central apparatus, C1b projection, cryo-electron tomography, subtomogram averaging, quantitative mass spectrometry

## Abstract

‘9+2’ motile cilia contain 9 doublet microtubules and a central apparatus (CA) composed of two singlet microtubules with associated projections. The CA plays crucial roles in regulating ciliary motility. Defects in CA assembly or function usually result in motility-impaired or paralyzed cilia, which in humans causes disease. Despite their importance, the protein composition and functions of most CA projections remain largely unknown. Here, we combined genetic approaches and quantitative proteomics with cryo-electron tomography and subtomogram averaging to compare the CA of wild-type *Chlamydomonas* with those of two CA mutants. Our results show that two conserved proteins, FAP42 and FAP246, are localized to the L-shaped C1b projection of the CA. We also identified another novel CA candidate protein, FAP413, which interacts with both FAP42 and FAP246. FAP42 is a large protein that forms the peripheral ‘beam’ of the C1b projection, and the FAP246-FAP413 subcomplex serves as the ‘bracket’ between the beam (FAP42) and the C1b ‘pillar’ that attaches the projection to the C1 microtubule. The FAP246-FAP413-FAP42 complex is essential for stable assembly of both the C1b and C1f projections, and loss of any of these proteins leads to ciliary motility defects. Our results provide insight into the subunit organization and 3D structure of the C1b projection, suggesting that the FAP246-FAP413-FAP42 subcomplex is part of a large interconnected CA-network that provides mechanical support and may play a role in mechano-signaling between the CA and radial spokes to regulate dynein activity and ciliary beating.

**Summary Statement:** The present work provides insight into the subunit organization and 3D structure of the C1b projection of CA and the mechanism by which it regulates dynein activity and ciliary beating.

## Introduction

Cilia and flagella (terms here used interchangeably) are highly conserved organelles in eukaryotes, where they play a variety of roles in cells, including motility, generating fluid flow, and sensing extracellular signals. A wide range of human diseases, collectively known as ciliopathies, are associated with cilia dysfunction, leading to symptoms including chronic respiratory infections, laterality abnormalities, and infertility (Afzelius, 2004; Braun and Hildebrandt, 2017; Brown and Witman, 2014). The core component of the motile cilium is the ‘9+2’ axoneme, which consists of nine outer doublet microtubules (DMTs) and a central apparatus (CA) composed of two singlet microtubules and associated projections. Each DMT consists of many copies of 96-nm-long units that repeat along the axoneme; each repeat includes major substructures such as outer and inner dynein arms (ODAs and IDAs), radial spokes (RSs), and nexin-dynein regulatory complexes (N-DRCs) (Fig. 1A). Together with the CA, these substructures produce and regulate the cilium’s motility, and distinguish motile cilia from non-motile cilia such as primary cilia (Dymek and Smith, 2007; Gui et al., 2019; Kikkawa, 2013; Lin and Nicastro, 2018; Loreng and Smith, 2017; Nicastro et al., 2006; Roberts et al., 2013; Viswanadha et al., 2017).

**Figure 1.**
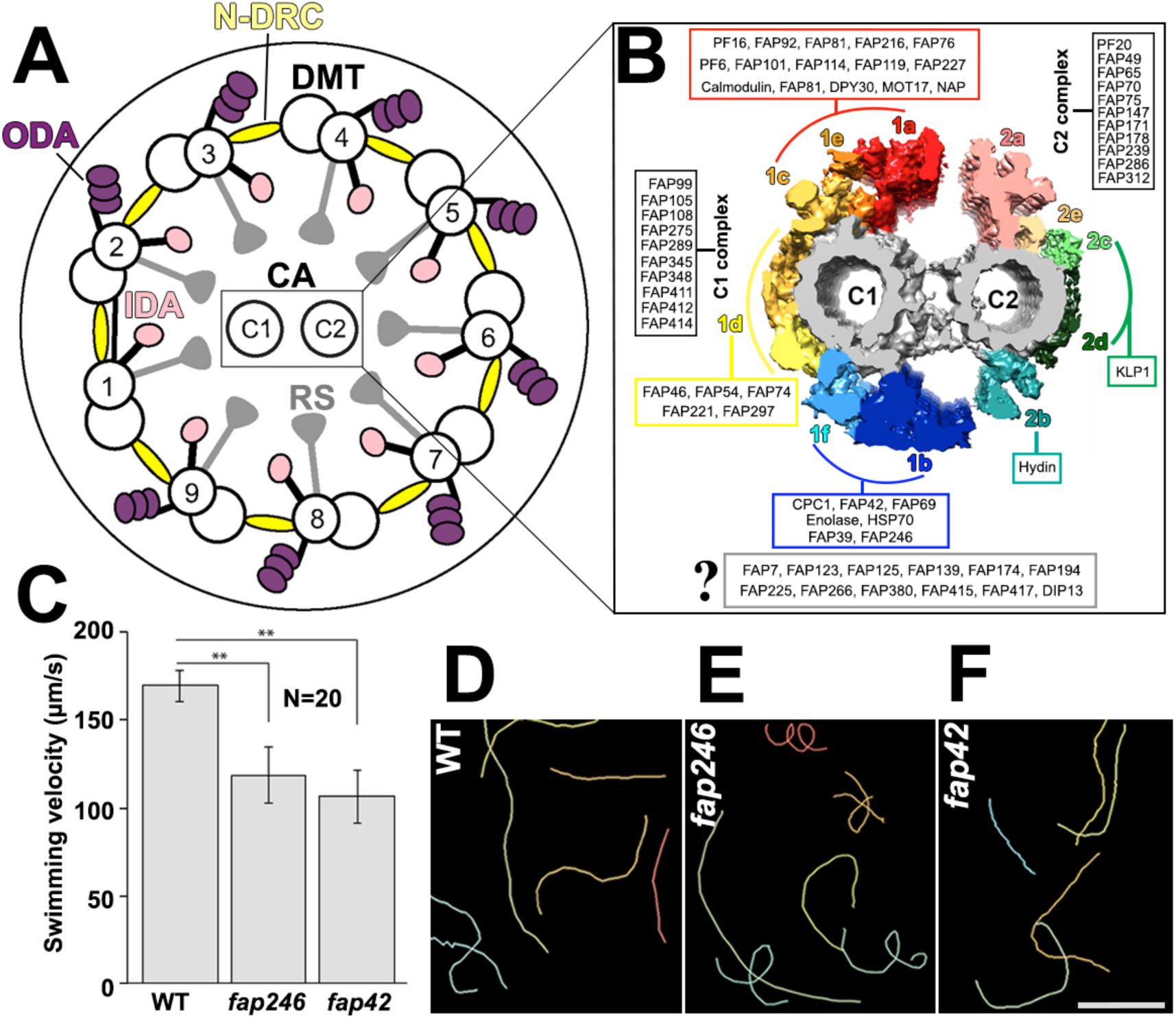
Known structure and proteome of the CA, and motility phenotypes of *Chlamydomonas* CA mutants. **A)** Schematic of a *Chlamydomonas* flagellum in cross-sectional view, showing the doublet microtubules with inner and outer dynein arms (IDA, ODA), the nexin-dynein regulatory complex (N-DRC), and the radial spokes (RS) that connect to the central apparatus (CA); cross-sections are viewed from proximal (flagellar base) in all figures, unless otherwise stated. **B)** Isosurface rendering of the averaged *Chlamydomonas* CA in cross-sectional view and predicted locations of CA proteins based on previous proteomic studies [modified after (Zhao et al., 2019)]. Naming and coloring of CA projections are adapted from (Carbajal-Gonzalez et al., 2013; Fu et al., 2019). **C-F)** Average swimming velocities (**C**) and swimming trajectories (**D-F**) of *Chlamydomonas* WT and two CA mutant cells, i.e. *fap246* and *fap42*. The error bars in (**C**) represent the standard deviations from 20 cells. Significant difference between WT and mutant strains as indicated (Student’s t test, P<0.01). In (**D-F**), each colored line represents the swimming path of one cell recorded for 2 s. Scale bar: 100 µm.

The CA is the largest ciliary regulatory complex. The two singlet microtubules, termed C1 and C2, are both canonical 13-protofilament microtubules; however, they differ in stability and associated projections. Recent cryo-electron tomography (cryo-ET) studies of the *Chlamydomonas reinhardtii* axoneme have identified at least 11 protein projections with a total molecular mass of > 14 MD that form 16- or 32-nm repeating units along the C1 and C2 microtubules (Carbajal-Gonzalez et al., 2013). Among the 11 projections, six (C1a-f) are associated with the C1 microtubule and five (C2a-e) with the C2 microtubule (Fig. 1B and Movie S1). The projections consist of many proteins that connect the C1 and C2 microtubules and/or extend toward the RS heads. Presumably, the CA regulates ciliary motility through the CA→RS→N-DRC→IDA signaling pathway (Wirschell, 2009); however, the molecular details are still elusive. The CA is essential for normal ciliary motility from *Chlamydomonas* to mammals, and mutations affecting the CA often lead to impaired or paralyzed cilia (Dutcher et al., 1984; Smith and Lefebvre, 1996; Smith and Yang, 2004; Witman et al., 1978). In mice and humans, defects in CA proteins are associated with the ciliopathy primary ciliary dyskinesia (PCD) (Loreng and Smith, 2017; Poprzeczko et al., 2019; Teves et al., 2016). For instance, defects in the C1d proteins FAP221 (also known as Pcdp1) or FAP54 lead to typical PCD symptoms in mice, including accumulation of mucus in the lungs, male infertility, and hydrocephalus (Lee et al., 2008; McKenzie et al., 2015). In humans, mutations in the CA protein PF20 (human Spag16L) can cause male infertility, and recessive mutations in the C2b protein Hydin cause PCD (Olbrich et al., 2012; Zhang et al., 2007). Therefore, it is important to establish a detailed understanding of CA structures and the mechanisms by which the CA regulates ciliary motility.

Despite the importance of the CA for ciliary motility, the specific locations of most of its proteins remain unknown, which makes CA the least understood axonemal structure to date. Several recent proteomic and structural studies have shed light on the protein composition and 3D-structure of the CA (Carbajal-Gonzalez et al., 2013; Dai et al., 2020; Fu et al., 2019; Zhao et al., 2019). For example, proteomic comparisons of CAs from mutant and wild-type (WT) *Chlamydomonas* identified 37 and 44 new candidate CA proteins, respectively (Dai et al., 2020; Zhao et al., 2019), of which many were assigned to the C1 or C2 microtubule and some to specific projections (Fig. 1B). Our recent cryo-ET study visualized the CA structure with up to 2.3 nm resolution and provided details about the molecular architecture of the PF16-dependent C1a-e-c supercomplex that contains at least 16 proteins (Fu et al., 2019). However, due to the high connectivity within and between CA projections, the assignment of CA proteins to certain projections based on biochemical and mass spectrometry (MS) data can lead to ambiguous or even erroneous conclusions. For instance, the two CA proteins FAP76 and FAP81 were unambiguously shown to be components of the C1a-e-c supercomplex by cryo-ET studies (Fu et al., 2019), but were mistakenly assigned to the C1d projection in a proteomic study (Dai et al., 2019). Given this caveat, it is important to complement MS studies with structural studies, e.g. using cryo-ET of mutants or tagged candidate proteins, to localize specific subunits within the CA.

The C1b projection is one of the largest and most complex CA projections, which appears to have a particularly strong affinity for RS heads (Nakazawa et al., 2014). Pioneering studies on this projection in *Chlamydomonas* showed that it physically interacts with the C2b projection of the C2 microtubule, and that the *cpc1* mutant, which swims slowly due to reduced flagellar beat frequency, lacks the entire C1b projection as assessed by conventional EM, suggesting that the C1b projection has a role in the control of beat frequency (Mitchell and Sale, 1999). Biochemical analyses found that the *cpc1* axonemes lacked three proteins that co-sedimented as part of a larger complex, termed the CPC1 complex, containing at least six proteins (Mitchell and Sale, 1999). Further analysis identified five of these proteins as CPC1, FAP42, FAP69, enolase and HSP70A (Fig. 1B) (Mitchell et al., 2005). The first cryo-ET study of the CA (Carbajal-Gonzalez et al., 2013) revealed that the C1b projection is more architecturally complex than previously known, and its estimated mass is nearly 2 MDa, i.e. twice the sum of the masses of the known C1b subunits (883 kDa). Cryo-ET also showed that C1b interacts with a previously undescribed projection, C1f, that is located between the C1b and C1d projections (Carbajal-Gonzalez et al., 2013). In our previous analyses of the CA proteome, we identified two novel CA candidate proteins, FAP39 and FAP246, that were assigned to the C1 microtubule and co-immunoprecipitated with known C1b proteins, indicating that they likely are C1b or C1f proteins. Axonemes of the mutant *fap246-1* had greatly reduced amounts of both FAP246 and another candidate CA protein, FAP413 (Zhao et al., 2019). However, none of these proteins have been localized precisely within the C1b or C1f projections.

Here, we integrated genetic and quantitative proteomic approaches with cryo-ET and subtomogram averaging to compare CAs from mutants for two highly conserved and predicted C1b proteins, FAP42 and FAP246, with the WT CA. The *Chlamydomonas fap42* and *fap246* mutants showed impaired motility compared to WT, indicating their importance in the regulation of ciliary beating. Our data allowed localization of FAP42, FAP246 and FAP413 within the C1b projection, as well as refined predictions for the locations of FAP39 and six additional CA proteins. Classification analyses of the CA structures of the mutants revealed that the FAP246-FAP413-FAP42 subcomplex is critical for the stability of neighboring structures and may play a role in mechano-signaling from the CA to the RS and dyneins to regulate ciliary beating.

## Results

### *Chlamydomonas fap246* and *fap42* mutants have motility defects

FAP246 is a 120-kDa protein that is predicted to contain a highly conserved leucine-rich repeat (LRR) domain, a papain-like domain, and an EF-hand domain (Fig. S1A). We first reexamined the *fap246* mutant strain that was partially characterized in our previous study (Zhao et al., 2019). Nearly all *fap246* cells were motile, but they swam ∼30% slower than WT cells (Fig. 1C), a reduction in swimming velocity slightly greater than we previously reported (possibly reflecting culture in liquid TAP medium here vs. modified M medium) and similar to that of the *cpc1* mutant (Mitchell and Sale, 1999). Compared to WT, many *fap246* mutant cells also had an altered swimming pattern in which cells frequently changed swimming direction, resulting in highly curved and spiraling swimming paths (c.f. Fig. 1D and E). Therefore, FAP246 is likely to have an important role in the control of flagellar motility.

FAP42 is a conserved ∼300-kDa protein that is predicted to have a papain-like domain and four guanylate kinase domains (Fig. S1C). We obtained a *fap42* mutant strain (CLiP ID: LMJ.RY0402.205930) from the CLiP library (Li et al., 2016) and, using PCR, confirmed insertion of the cassette into the *FAP42* gene in a genetically homogenous clone (Fig. S1F). Compared to WT cells, *fap42* mutant cells swam ∼37% slower (Fig. 1C). Cells of *fap42* swam slightly slower than *fap246* cells, but their swimming paths were less curved (Fig. 1F). The results suggest that FAP42 also is important for the regulation of flagellar motility.

### Proteomics studies of *fap246* and *fap42* axonemes

We used quantitative MS to compare the axonemal proteomes of WT, *fap42* and *fap24*6 axonemes. Label-free quantitation of proteins across samples was performed using SINQ normalized spectral index Software (Trudgian et al., 2011) (Table 1). We included the recently identified novel candidate CA proteins (Dai et al., 2020; Zhao et al., 2019) in our *Chlamydomonas reinhardtii* protein database during the peptide search (Table S1). The MS/MS results confirmed that the protein level of FAP246 is significantly reduced in the *fap246* axoneme compared to WT (*fap246*/WT ratio = 0.05). In the *fap246* axoneme, MS detected 8 exclusive unique FAP246 peptides compared with 46 in the WT samples (Table 1). The 8 exclusive unique FAP246 peptides that were identified originated from the N-terminus of the protein, suggesting that a small amount of C-terminally truncated FAP246 was assembled into the *fap246* axonemes (Table S2). The only other protein that was significantly reduced in the *fap246* axoneme was FAP413 (*fap246*/WT ratio = 0.02). Only 2 exclusive unique FAP413 peptides were detected in the *fap246* axoneme compared with 44 in that of WT (Table 1 and S2), indicating that very little FAP413 is assembled into *fap246* axonemes. FAP413 is a 210-kDa protein that is predicted to have a large WD40-repeat domain at the C-terminus with at least 11 WD40 repeats (Fig. S1E). The results indicate that FAP246 and FAP413 are likely in the same CA substructure, and that FAP413 is dependent on FAP246 for its assembly into the axoneme.

**Table 1.** MS/MS analysis of the axonemes of fap246 and fap42 mutants — CA proteins previously or here confirmed to be localized to a specific CA projection.

| Protein | CA | Phytozome gene | MW (kDa) | No. of Unique Peptides |  |  | Quantification Ratio<sup>a</sup> |  |
| --- | --- | --- | --- | --- | --- | --- | --- | --- |
|  | projection | number |  | WT | *fap246* | *fap42* | *fap246*/WT | *fap42*/WT |
| CA proteins<sup>b</sup> |  |  |  |  |  |  |  |  |
| FAP246 | C1b | Cre14.g618750 | 120 | 46 | 8 | 36 | 0.05 | 0.75 |
| FAP42 | C1b | Cre12.g519950 | 262 | 135 | 137 | 29 | 1.02 | 0.04 |
| FAP413 | C1b | Cre08.g364000 | 215 | 44 | 2 | 10 | 0.01 | 0.03 |
| FAP39 | C1b | Cre02.g145100 | 127 | 29 | 27 | 16 | 0.83 | 0.33 |
| FAP69 | C1b | Cre03.g168200 | 110 | 45 | 45 | 45 | 1.03 | 1.03 |
| CPC1 | C1b | Cre03.g183200 | 200 | 118 | 112 | 119 | 0.98 | 1.18 |
| Enolase | C1b | Cre12.g513200 | 52 | 36 | 35 | 35 | 0.85 | 0.64 |
| HSP70A | C1b | Cre08.g372100 | 71 | 47 | 44 | 40 | 0.79 | 0.40 |
| FAP54 | C1d | Cre12.g518550 | 319 | 121 | 127 | 125 | 1.05 | 1.13 |
| FAP46 | C1d | Cre10.g420800 | 291 | 123 | 124 | 126 | 1.00 | 1.10 |
| FAP221 | C1d | Cre11.g476376 | 71 | 25 | 24 | 24 | 0.95 | 1.04 |
| FAP297 | C1d | Cre01.g029350 | 74 | 41 | 41 | 43 | 0.92 | 0.96 |
| FAP74 | C1d | Cre06.g271150 | 200 | 64 | 66 | 67 | 1.07 | 1.17 |
| PF6 | C1a | Cre10.g434400 | 237 | 100 | 104 | 107 | 1.11 | 1.05 |
| PF16 | C1a | Cre09.g394251 | 50 | 26 | 25 | 24 | 0.97 | 0.91 |
| FAP92 | C1a | Cre13.g562250 | 150 | 60 | 59 | 66 | 1.16 | 1.06 |
| FAP101 | C1a | Cre02.g112100 | 86 | 27 | 27 | 27 | 1.01 | 1.04 |
| FAP119 | C1a | Cre06.g256450 | 34 | 12 | 12 | 12 | 1.13 | 0.81 |
| FAP227 | C1a | Cre17.g729850 | 17 | 17 | 17 | 16 | 1.01 | 0.96 |
| FAP114 | C1a | Cre09.g389282 | 32 | 15 | 14 | 13 | 0.91 | 0.79 |
| **FAP81** | C1e | Cre06.g296850 | 172 | 68 | 68 | 68 | 1.07 | 1.06 |
| **FAP216** | C1c | Cre12.g497200 | 79 | 33 | 31 | 31 | 0.74 | 0.89 |
| **FAP76** | C1c | Cre09.g387689 | 162 | 76 | 82 | 76 | 1.11 | 1.02 |
| **Hydin** | C2b | Cre01.g025400 | 528 | 209 | 217 | 212 | 1.06 | 1.16 |

<sup>a</sup>Missing or significantly reduced unique peptide numbers (less than half of WT) or ratios (<0.2) are highlighted in bold.
<sup>b</sup>FAP246, FAP413 and FAP39 are novel proteins that are predicted to be CA proteins by our recent proteomic study (Zhao et al., 2019).

The MS analyses of *fap42* axonemes revealed that the protein level of FAP42 is significantly reduced in the mutant axoneme compared to WT (*fap42*/WT ratio = 0.04) (Table 1). In the *fap42* axoneme, MS detected 29 exclusive unique FAP42 peptides compared with 135 exclusive unique peptides in the WT sample (Table 1). These peptides originated from throughout the FAP42 sequence (Table S3), suggesting that a small amount of full-length FAP42 is assembled into the *fap42* mutant axoneme. The only other protein that was significantly reduced in the *fap42* axoneme was FAP413 (*fap42*/WT ratio = 0.03). 10 exclusive unique FAP413 peptides were detected in the MS analyses of *fap42* axonemes compared with 44 in that of WT (Table 1 and S3), indicating that very little FAP413 is assembled into the mutant axoneme. The results suggest that FAP413 and FAP42 assemble into the same complex, and that FAP42 is essential for proper assembly of FAP413 into the axoneme. The level of FAP246 in *fap42* axonemes was only slightly decreased (*fap42*/WT ratio = 0.75) (Table 1). The fact that FAP413 was significantly decreased in the MS analyses of both *fap42* and *fap246* axonemes suggests that it interacts with both FAP42 and FAP246.

Several other proteins have been predicted to be in the C1b projection, i.e. CPC1, FAP39, FAP69, the chaperone protein HSP70A, the glycolytic enzyme enolase, and, with less confidence, FAP174 and FAP380 (Fig. 1B) (Mitchell et al., 2005; Zhang and Mitchell, 2004; Zhao et al., 2019). Our MS/MS analyses showed that the protein level for none of these proteins was significantly affected in *fap246* axonemes. In the MS/MS analyses of *fap42* axonemes, the protein levels of FAP69 and CPC1 were not affected, but the levels of HSP70A, enolase, FAP39, and FAP380 were reduced to 40%, 64%, 33% and 36% of WT level, respectively (Table 1 and S1). HSP70A and enolase are known to locate not only to the C1b projection but elsewhere in the axoneme (Mitchell et al., 2005; Zhao et al., 2019), so that the amounts remaining in *fap42* axonemes could represent the non-C1b pool of these proteins. None of the known proteins in the C1-a-e-c supercomplex or C1d projection were significantly affected (Table 1 and S1).

### FAP246 is localized to the C1b projection and forms a complex with FAP413

We next carried out cryo-ET and subtomogram averaging to investigate the CA structure of *fap246* and compare it with the WT CA. Our subtomogram averages revealed the CA structures of WT and *fap246* axonemes with 2.5-nm and 2.2-nm resolution, respectively (Fourier shell correlation 0.5 criterion) (Fig. S1G), similar to what we reported previously (Fu et al., 2019). In WT, C1b is one of the largest CA projections (∼1.3 MDa) that has a roughly triangular shape in cross-sectional view. We named the three sides of the triangle: the 17 nm long “pillar” that projects from C1 protofilament #9 outwards, in the periphery a long perpendicular “beam”, and the “bracket” that connects diagonally between pillar and beam (Fig. 2A-D and the inset). The bracket density consists of two roughly globular domains of which the slightly more proximal-central one (the “inner bracket”) connects to the pillar, and the more distal-peripheral one (the “outer bracket”) links to the beam (Fig. 2 B and D). The beam structure, but not the bracket, extends longitudinally and connects with the beam structures of the neighboring repeats along the axoneme length, and also forms a small connection with the C2b projection (Movie S1). All C1b structures have a 16 nm periodicity and form multiple connections to neighboring CA structures.

**Figure 2.**
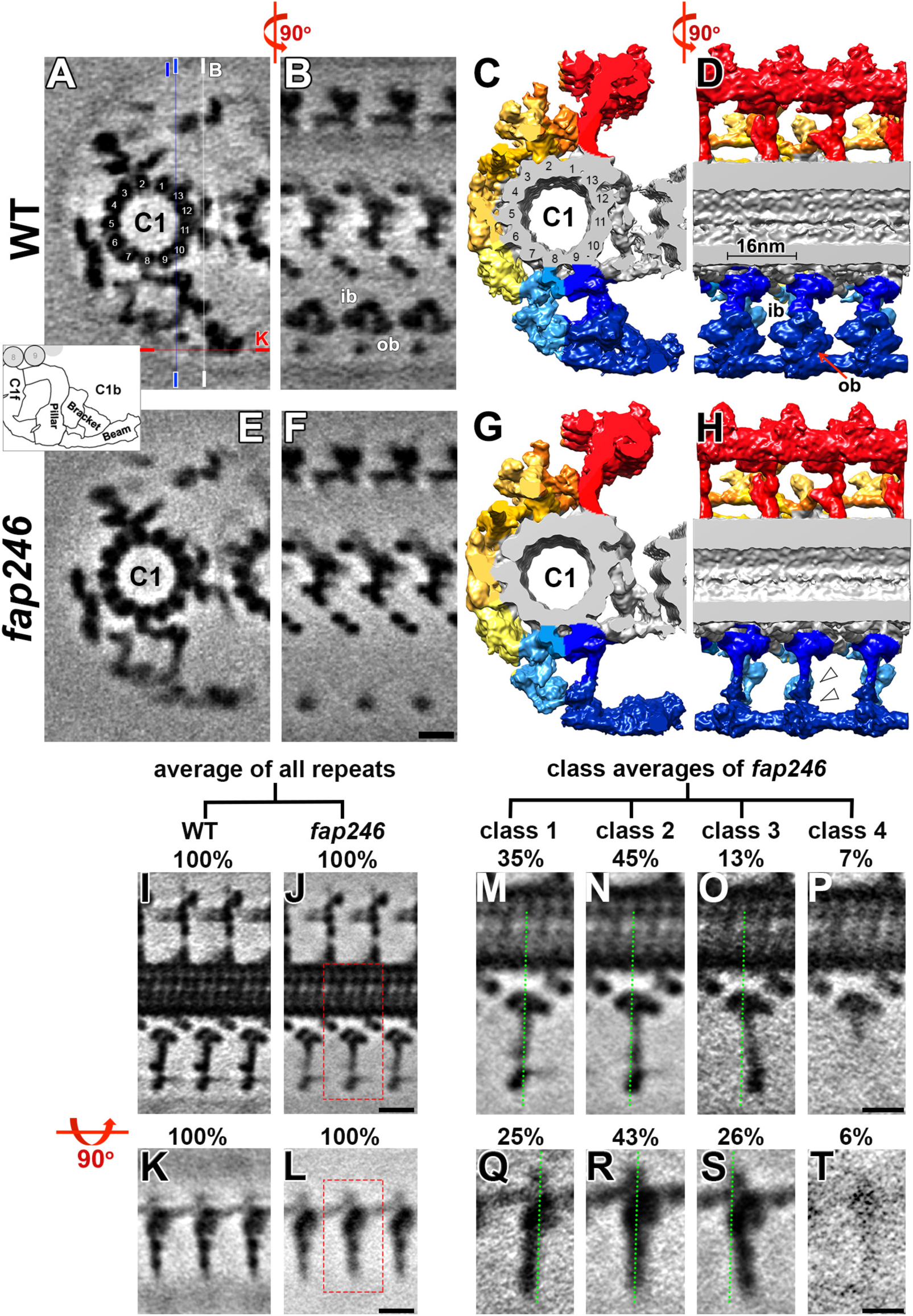
Cryo-ET, subtomogram averaging and classification analyses of the *fap246* CA revealed defects in the C1b projection and that FAP246 plays a role in stabilizing the C1b structure. **A-H)** Tomographic slices (**A, B, E** and **F**) and isosurface renderings (**C, D, G** and **H**) of the averaged CA repeats of WT (**A-D**) and *fap246* (**E-H**) in cross-sectional (**A**, **C**, **E** and **G**) and longitudinal views (**B**, **D**, **F** and **H**) show that the C1b inner and outer bracket (ib, ob) densities are present in WT, but missing in *fap246* (white arrowheads). The thin lines in (**A)** indicate the locations for the slices shown in (**B** and **F**, white line), (**I** and **J**, blue line) and (**K** and **L**, red line). In longitudinal slices proximal is on the left, unless otherwise stated. *Inset*, schematic of the C1b projection shows pillar, bracket and beam regions. **I-T)** Classification analyses revealed structural heterogeneity in *fap246* but not the WT C1b projection. Tomographic slices (longitudinal) of the averages that included all CA repeats showed that the pillar (**I** and **J**) and the beam (**K** and **L**) densities were blurred in the *fap246* mutant (**J** and **L**) compared to WT (**I** and K). Classification analyses on these regions revealed distinct classes (percentages of repeats are indicated for each class) for both the pillar (4 classes) and the beam (4 classes). The regions shown in (**M-P**) and (**Q-T**) are indicated by the red dotted boxes in (**J**) and (**L**), respectively. The green dotted lines serve as references to show the positional differences between classes. Scale bars: 10 nm (in F valid for A, B, E and F; in J valid for I and J; in L valid for K and L); 5 nm (P valid for M-P; T valid for Q-T).

In *fap246,* the averaged CA repeats lacked the entire bracket density of the C1b projection, confirming that FAP246 is a C1b protein (Figs. 2E-H; S2). The molecular weight (MW) of the FAP246-dependent density was estimated to be ∼350 kDa based on the volume difference between the mutant and WT CA average (for details see Methods). This suggested that the bracket either contains three copies of FAP246 (120 kDa) or additional C1b subunit(s). The only protein besides FAP246 that was significantly reduced in the MS analyses of *fap246* axonemes was FAP413 (210 kDa). The MWs of the inner and outer bracket (Fig. 2B and D, ib and ob) domains were estimated to be 150 kDa and 180 kDa in WT, respectively. This suggested that FAP246 and FAP413 form a subcomplex, with FAP246 localizing to the inner and FAP413 to the slightly larger outer bracket domain, and that the stable assembly of FAP413 into axonemes required FAP246. Previously, FAP413 was predicted to be a C1 protein, but could not be assigned to a specific projection (Zhao et al., 2019).

Compared to the WT CA, the density of both the pillar and the beam appeared to be weakened and blurred in the *fap246* CA structure (Fig. 2I-L). This blurring suggests either partial reduction of the structures (i.e. not all repeats contain them) or positional flexibility of them. Therefore, we applied automatic image classification analyses with a mask that focused on the blurred densities (Heumann et al., 2011). Classification of the WT CA structure in this region revealed one structurally homogenous class. In contrast, the C1b structure of *fap246* was separated into four structurally different classes (Fig. 2M-P, Movie S2): in 45% the pillar was positioned in the center like in WT (Fig. 2N, class 2), in 35% the pillar tilted towards the axoneme base (Fig. 2M, class 1), in 13% the pillar tilted towards the axoneme tip (Fig. 2O, class 3), and in 7% of the repeats the distal half of the pillar was missing (Fig. 2P, class 4). The 3D classification analyses on the blurred beam density also revealed four classes with comparable distribution as the pillar classes, which is consistent with the pillar and beam forming a unit that moves together (Fig. 2Q-T). Specifically, in 43% the beam density was positioned in the center like in WT (Fig. 2R, class 2), in 25% the beam density tilted towards the axoneme base (Fig. 2Q, class 1), in 26% the beam tilted towards the axoneme base (Fig. 2S, class 3), and in 6% of the repeats the beam density was missing (Fig. 2T, class 4). These results showed both positional flexibility and a small reduction of the pillar and beam densities in *fap246*, strongly suggesting that the FAP246-FAP413 bracket plays a role in stabilizing the neighboring C1b pillar and beam.

### FAP42 forms the beam of the C1b projection

FAP42 was one of the 5 originally identified C1b proteins in the CPC1 complex (Mitchell et al., 2005), but so far it has not been localized within the C1b structure. Using cryo-ET and subtomogram averaging, we visualized the *fap42* CA with 2.5 nm resolution (FSC 0.5 criterion) (Fig. S1G). Compared to the WT CA, the averaged *fap42* CA repeats lacked the entire C1b beam and the outer bracket density that usually connects to the beam (Figs. 3 and S2). The mass of the entire FAP42-dependent density was estimated to be ∼480 kDa, i.e. ∼300 kDa for the beam and ∼180 kDa for the outer bracket structure. The only two proteins that were almost completely reduced in the MS analyses of *fap42* axonemes were FAP42 and FAP413 (Table 1). This suggests that FAP42 and FAP413 form a subcomplex, with FAP42 (260 kDa) forming (most of) the beam structure, and FAP413 localizing to the outer bracket domain, in good agreement with our results for the *fap246* mutant. Similar to FAP246, FAP42 is also required for stable assembly of FAP413 into the CA. The FAP42-FAP413 subcomplex is connected to several neighboring densities, i.e. FAP413 (outer bracket) connects to FAP246 (inner bracket), while FAP42 (the beam) connects with the peripheral tip of the C1b pillar, with the distal tips of the beams in the neighboring repeats, and with apposing structures of the C2b projection (Fig. 3A, D and E). Our MS analyses showed that the protein level of FAP413 is significantly reduced in both *fap42* and *fap246* axonemes, which is consistent with it being the shared missing density in the averaged CA repeats of *fap42* and *fap246* axonemes (Fig. 2 and 3, Table 1). Thus our results revealed that FAP42 forms most of the C1b beam, FAP413 the outer bracket domain and FAP246 the inner bracket domain.

**Figure 3.**
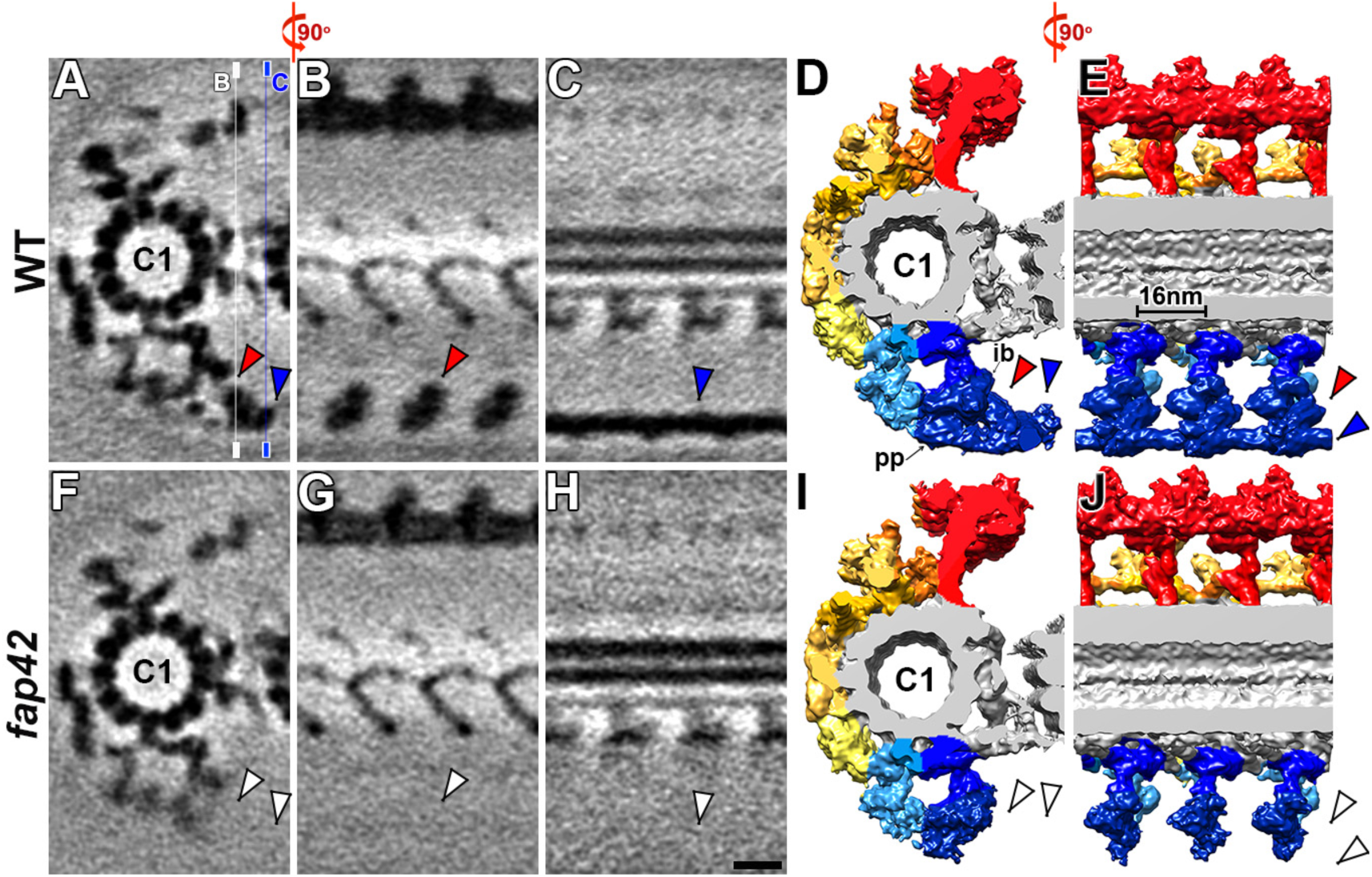
Cryo-ET and subtomogram averaging localize FAP42 to the C1b projection. **A-J)** Tomographic slices (**A-C** and **F-H**) and isosurface renderings (**D, E, I** and **J**) of the averaged CA repeats of WT (**A-E**) and *fap42* (**F-J**) in cross-sectional (**A**, **D**, **F** and **I**) and longitudinal views (**B**, **C**, **E, G, H** and **J**) show that the structures that are present in the wild-type CA repeats (red and blue arrowheads) but missing in those of *fap42* (white arrowheads). The thin lines in (**A)** indicate the locations for the slices shown in (**B** and **G**, white line) and (**C** and **H**, blue line). Other labels: ib, inner bracket density; pp, peripheral part of the pillar. Scale bar: 10 nm (in H valid for all EM images).

Compared to the WT CA, several structures that interact with the FAP42-dependent complex were weakened or blurred in *fap42* (Fig. 4A-H and A’), including the FAP246 inner bracket (compare Fig. 4A, B with 4E and F, red arrowheads), the peripheral part of the C1b pillar (compare Fig. 4A, C with E and G, blue arrowheads), and the C1f projection that connects to the pillar structure (compare Fig. 4A, D and E and H, green arrowheads). We applied automatic 3D image classification analyses with masks focusing on the above weakened C1b and C1f densities (Heumann et al., 2011). In all cases, the classification analyses of WT CA repeats showed one structurally homogenous class. In contrast, for the two weakened C1b densities (FAP246 and the peripheral part of the C1b pillar), two distinct classes were identified for *fap42*: in class 1 (54%) both densities were present with full occupancy like in WT (Fig. 4I-K), whereas they were missing in the class 2 (46%) (Fig. 4L-N). For the C1f projection, three classes were identified for *fap42*: in class 1 (50%) the C1f density connecting with the C1b pillar was missing (Fig. 4O and P), in classes 2 and 3 (26 and 24%) the same C1f density was present with full occupancy, but the C1f structures showed positional flexibility (Movie S3), i.e. in class 2 it was tilted proximal along the C1 microtubule (Fig. 4Q and R) compared to the WT-like class 3 (Fig. 4S and T). The 3D classification analyses revealed that loss of the FAP42-FAP413 subcomplex resulted in both reduced occupancy and positional flexibility of the neighboring C1b pillar and C1f projection. Therefore, FAP42 is essential for the stable assembly and positioning of neighboring structures in the C1b and C1f projections.

**Figure 4.**
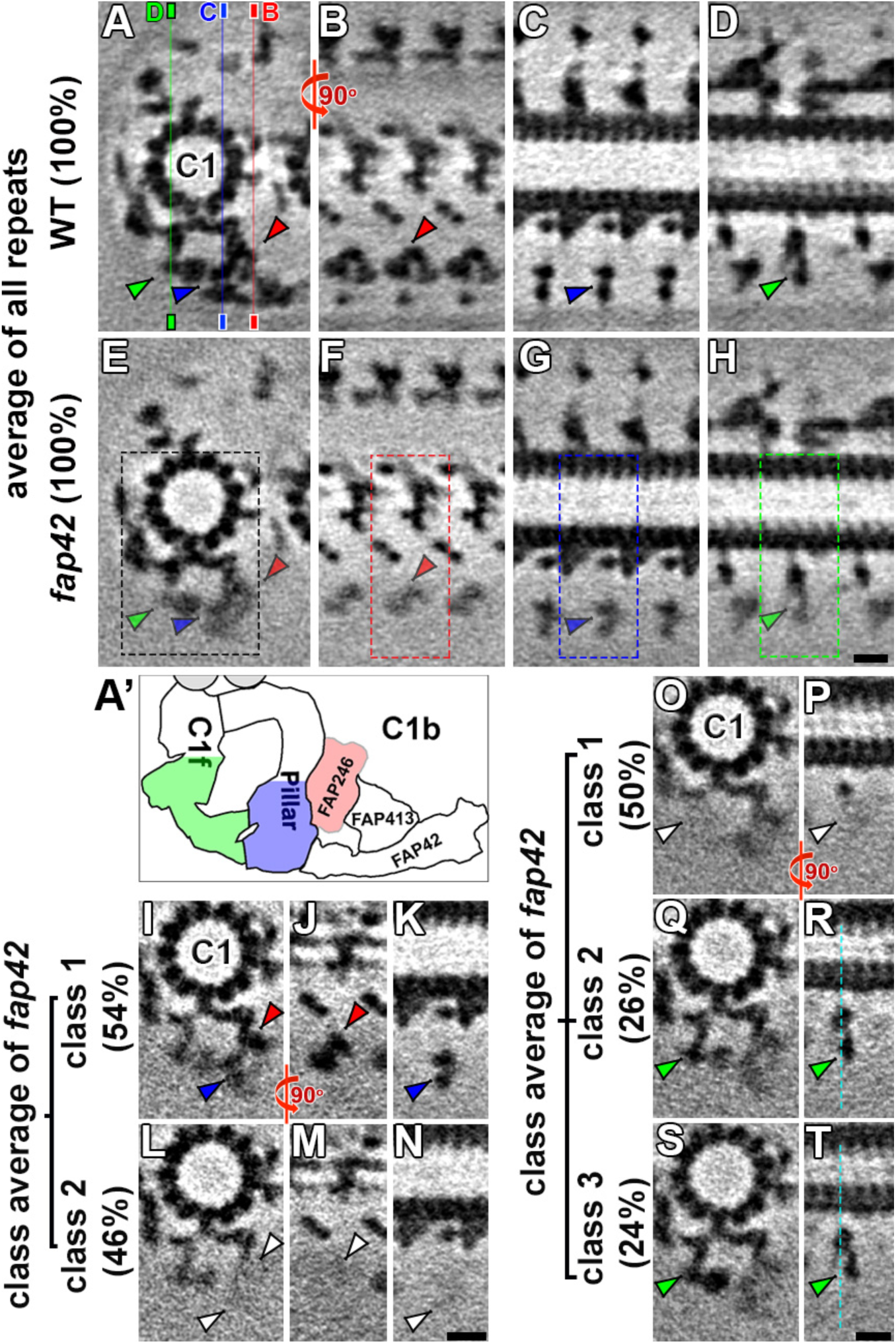
Classification analyses revealed structural heterogeneity for the C1b and C1f projections in *fap42* but not for WT. **A-H)** Tomographic slices of averaged CA repeats of WT (**A**-**D**) and *fap42* (**E**-**H**) axonemes viewed in cross-sectional (**A** and **E**) and longitudinal orientations (**B**-**D** and **F**-**H**). Several C1b and C1f densities, including FAP246 (red arrowheads), the peripheral part of the pillar (blue arrowheads) and the peripheral part of C1f (green arrowheads) were blurred in the averages of all CA repeats of *fap42* axonemes (**E-H**) compared to that of WT (**A-D**). The thin lines in (**A)** indicate the locations for the slices shown in (**B** and **F**, red line), (**C** and **G**, blue line) and (**D** and **H**, green line). (**A’**) Schematic of the C1b (with pillar, FAP246-FAP413 bracket and FAP42 beam) and the C1f projections. The red, blue and green colored regions indicate regions that were blurred in the average of all *fap42* CA repeats (**A**-**H**). **I-T)** Classification analyses on these regions revealed distinct classes (percentages of repeats are indicated for each class) for the peripheral regions of both the C1b (2 classes) and C1f projections (3 classes); cross-sectional (**I**, **L**, **O**, **Q** and **S**) and longitudinal views (**J**, **K**, **M**, **N**, **P**, **R** and **T**). The regions shown in (**I, L, O-S**), (**J** and **M**), (**K** and **N**), and (**P**, **R** and **T**) are indicated by the dotted boxes in (**E**), **(F), (G)** and (**H**), respectively. The red, blue and green arrowheads indicate presence of the densities in classes, whereas the white arrowheads indicate missing densities. The green dotted lines in (**R** and **T**) serve as references to show the positional differences of structures in different classes. Scale bars: 10 nm (in H valid for A-H; in N valid for I-N; in T valid for O-T).

### C1b pillar interacts with several structures in the C1b and C1f projections

Our cryo-ET results showed that the C1b “pillar” is highly connected with neighboring CA structures through six interfaces (Fig. 2A and C; and Fig. 5, dark blue). We observed two connections from the pillar within the C1b projection, i.e. to the FAP42 beam (Fig. 5, green), and to the FAP246 inner bracket domain (Fig. 5D, magenta colored density). Additional interfaces of the C1b pillar are: the attachment site to protofilament 9 of the C1-microtubule, two connections with the C1f projection (Fig. 5D, cyan), and a connection to part of the CA bridge density that links the C1 and C2 microtubules (Fig. 5D, gray).

**Figure 5.**
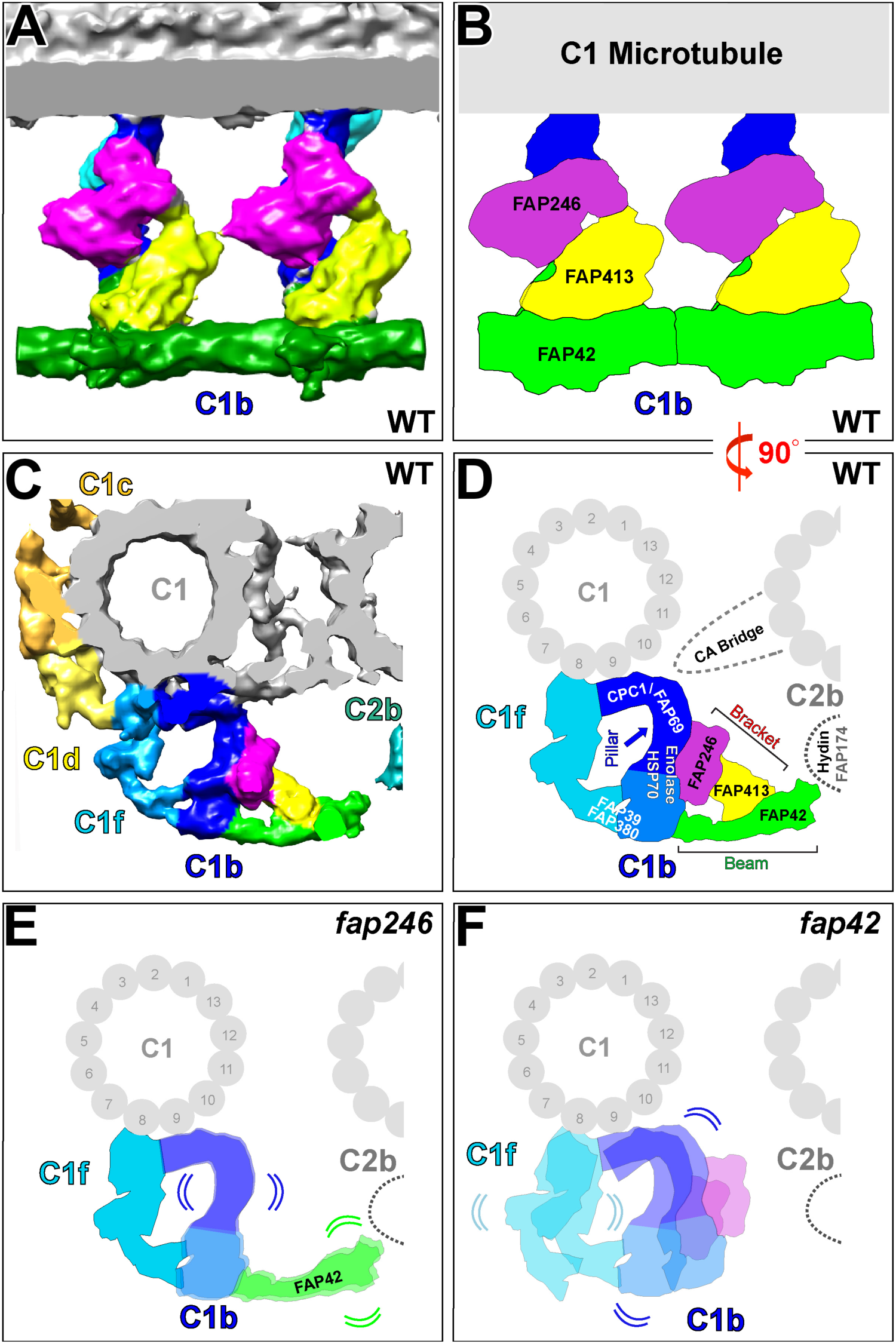
Summary model depicting the structural organization of the C1b projection, and showing the effects of partial loss of the FAP246-FAP413-FAP42 subcomplex on the CA structure. **A-D)** The isosurface renderings (**A** and **C**) and schematic drawings (**B** and **D**) show the 3D structure of the averaged *Chlamydomonas* WT CA repeat viewed in longitudinal orientation (**A** and **B,** two 16 nm repeats are shown) and cross-sectional orientation (**C** and **D**). The views highlight the subunit organization of the C1b projection, including FAP246 (magenta), FAP413 (yellow), and FAP42 (green). The predicted localizations of neighboring candidate proteins in the C1b, C1f and C2b projections are indicated. **E-F)** Schematic drawings of the averaged 16 nm CA repeat of *fap246* (**E**) and *fap42* (**F**) in cross-sectional view showing the observed structural defects in the mutant CAs; compared to WT, the densities of FAP246 and FAP413 are missing in *fap246*, and the pillar (blue) and the beam densities show positional flexibility. In *fap42*, the densities of FAP42 and FAP413 are missing, and neighboring C1b and C1f densities show positional flexibility, as well as partial reduction.

The protein composition of the pillar is unknown and cannot be precisely dissected using the CA mutants studied here, because the pillar structure remained mostly present in both mutants. However, previous studies predicted five additional C1b proteins, i.e. CPC1 (205 kDa), FAP69 (114 kDa), enolase (51 kDa), HSP70A (71 kDa), FAP39 (127 kDa) (Mitchell et al., 2005; Zhao et al., 2019), that we did not find in the C1b beam and bracket structures. Therefore, the C1b pillar likely contains (most of) these proteins, which in sum would be slightly larger (∼550 kDa) than the estimated total molecular weight of ∼450 kDa for the pillar based on the cryo-ET averages.

## Discussion

### Protein composition and structural organization of the C1b projection

The CA is highly conserved and plays essential roles in regulating ciliary beating, likely through the [CA→RS→IDA & N-DRC→dyneins] signaling pathway (Wirschell, 2009). Single protein mutations often cause CA defects that result in impaired or paralyzed cilia, and are associated with various ciliopathies (Loreng and Smith, 2017; Poprzeczko et al., 2019; Teves et al., 2016). Our previous CA study combined structural and biochemical data, and revealed a 2 MDa C1-a-e-c supercomplex (Fu et al., 2019). We identified several key proteins in the C1a-e-c supercomplex, including PF16, FAP76, FAP81, FAP92, and FAP216, and precisely localized them by cryo-ET (Fu et al., 2019). Our study also highlighted that the structural data were often critical for guiding the interpretation of biochemical data; e.g. FAP81 was found to immunoprecipitate with C1a and C1e proteins, but it could not be definitively assigned to either of these projections (Zhao et al., 2019); in contrast to these biochemical results though, our cryo-ET study revealed that FAP81 localized to the C1c projection within the interconnected C1-a-e-c supercomplex (Fu et al., 2019). For most of the remaining CA projections, subunits have so far only been characterized biochemically, e.g. for the C1d projection (FAP46, FAP54, FAP74, FAP221, and FAP297), and the C2 projections (Brown et al., 2012; DiPetrillo and Smith, 2010; DiPetrillo and Smith, 2011; Lechtreck and Witman, 2007; Zhao et al., 2019).

The protein composition of the C1b projection has previously been predicted based on co-sedimentation in sucrose gradients, CA mutant proteomics and coimmunoprecipitation. The C1b candidate proteins included CPC1, FAP39, FAP42, FAP69, FAP246, the chaperone protein HSP70A, the glycolytic enzyme enolase, and - with less confidence – FAP174 and FAP380 (Dai et al., 2020; Mitchell et al., 2005; Zhang and Mitchell, 2004; Zhao et al., 2019). In the present study, we compared axonemes of WT and two C1b mutants, *fap42* and *fap246,* using a combination of quantitative proteomics and cryo-ET imaging. We localized FAP42 and FAP246 within the C1b projection, and identified and localized a new C1b subunit, FAP413; previously, FAP413 was thought to associate with the C1 microtubule, but could not be assigned to a specific CA projection (Zhao et al., 2019). Finally, we described the structural organization of the subcomplex formed by these 3 subunits (Fig. 5 and Movie S4).

The C1b projection can be divided into three major structural parts (Fig. 5 and Movie S4): 1) FAP42 (Fig. 5, green) forms repeating beams at the interface between the CA and RS heads; 2) the pillar (Fig. 5, dark blue) attaches the C1b protection to protofilament 9 of the C1 microtubule and is a major interaction hub with five additional interfaces with neighboring CA structures, including the C1f projection and C1-C2 bridge; and 3) FAP413 and FAP246 (Fig. 5, yellow and magenta, Movie S4) form a bracket between the pillar and beam; specifically, FAP413 forms the outer C1b bracket density that interacts with the beam, and FAP246 forms the inner bracket density that interacts with the pillar. We named the FAP246-FAP413 complex “bracket” because the C1b projection architecture resembles a pillar-beam construction for roofs, and our 3D-classification analyses revealed that disruption of the bracket structure in both *fap246* and *fap42* mutants resulted in structural destabilization (positional flexibility and reduction) of remaining C1b and C1f structures. All these structures have a 16-nm periodicity along the C1-microtuble, which is the predominant CA periodicity. A few other CA structures, such as features in the C1d projection, repeat with 32 nm instead of 16 nm periodicity (Carbajal-Gonzalez et al., 2013).

A previous study showed that FAP42 co-sediments with CPC1 (205 kDa), FAP69 (114 kDa), enolase (51 kDa) and HSP70A (71 kDa) in sucrose gradients (Mitchell et al., 2005) and that FAP39 co-immunoprecipiated with FAP246-HA (Zhao et al., 2019). These five previously identified C1b proteins were not completely missing in the MS of the here studied mutants of the C1b beam-bracket (FAP246-FAP413-FAP42) subcomplex (Table 1), suggesting that these proteins might instead form the C1b pillar and part of the C1f projection. This would be consistent with our size estimation for the pillar of ∼450 kDa. Previous classical electron microscopy studies observed that the entire C1b projection was missing from *Chlamydomonas cpc1* mutant flagella (Mitchell and Sale, 1999; Zhang and Mitchell, 2004). Therefore, CPC1 likely is located at the base of the C1b pillar and serves as a scaffold/docking protein that stabilizes the attachment of the C1b projection to the C1 microtubule, possibly by interacting also on two sides with the C1f projection and the C1-C2 bridge, respectively (Fig. 5). In contrast to CPC1 and FAP69 that were completely unaffected in our MS/MS analyses of *fap42* and *fap*246 axonemes, enolase, HSP70A, and FAP39 were reduced to between 33-64% of WT level in *fap42* and between 79-85% in *fap*246 (Table 1). This suggests that the latter proteins localize to the peripheral half of the C1b pillar (Fig. 5) that connects to FAP42 and FAP246, and was structurally missing in ∼46% of the *fap42* repeats (Fig. 4I-N, class 2) and in 7% of the *fap*246 repeats (Fig. 2, class 4). The estimated size of the missing pillar region is ∼450 kDa, slightly smaller than the sum of the five proteins (CPC1, FAP39, FAP69, HSP70, enolase) predicted to be pillar proteins. However, the borders between the C1b pillar and C1f projection were assigned based on previous classical EM studies of the *cpc1* mutant (Carbajal-Gonzalez et al., 2013).

Compared to cryo-ET, classical EM images have lower resolution, so that part of the C1f projection could actually belong to the C1b projection (Fig. 5), which would be consistent with the finding that part of the C1f projection also showed missing density in ∼50% of the repeats, similar to the peripheral pillar density.

To date, no C1f proteins have been identified. Our present proteomic analyses of *fap42* axonemes revealed that another candidate CA protein, FAP380 (20 kDa) (Zhao et al., 2019), was reduced to 36% of WT level (Table S1). Given the partial loss of the C1b pillar and C1f projection of *fap42*, FAP380 may be a subunit of the C1b pillar or C1f projection. None of the known proteins of the C1-a-e-c supercomplex or the C1d projection were decreased significantly in the C1b mutants studied here (Table 1).

The candidate CA protein FAP174 (10 kDa), which was previously assigned to C2 (Rao et al., 2016), was specifically enriched in one of the FAP246-HA coimmunoprecipitation replicates in Zhao et al.; therefore, FAP174 was predicted to be - with less confidence – either a C1b or C2b protein (Zhao et al., 2019). The C2b projection was not structurally affected in either of the mutants studied here, and FAP174 was not reduced in our MS of the mutants. Therefore, our results would be more consistent with FAP174 being a C2b, rather than a C1b/f, subunit (Fig. 5).

### C1b proteins may be involved in nucleotide cycles

FAP246 is a conserved protein that contains a highly conserved LRR domain and guanylate kinase domain (Fig. S1A). The mammalian homolog of FAP246, LRGUK-1 (NP_653249) (Fig. S1B), was previously localized to the acrosome-acroplaxoneme-manchette-tail network in mouse sperm (Liu et al., 2015). FAP42 is a conserved protein that contains four guanylate kinase domains (Fig. S1C). The human homolog of FAP42, guanylate kinase isoform b (GUK1; NP_000849.1) (Fig. S1D), catalyzes the transfer of a phosphate group from ATP to guanosine monophosphate (GMP) to eventually form guanosine diphosphate (GDP), and is a potential target for cancer chemotherapy (Khan et al., 2019). Guanylate kinase activity – possibly provided by both FAP42 and FAP246 - has not been studied in cilia, but maintenance of flagellar GTP concentrations may be important for both guanylate cyclase activity and tubulin polymerization (Mitchell et al., 2005). CPC1 contains an adenylate kinase domain, indicating that it might be involved in maintaining stable intraciliary ATP levels during ciliary beating (Zhang and Mitchell, 2004).

All studied C1b mutants, i.e. the previously described *cpc1* (Zhang and Mitchell, 2004) and the here characterized *fap246* and *fap42* (Fig. 1), showed similarly mild motility defects with ciliary beat frequency reduced to 60-70% that of WT, in contrast to mutations of some other CA proteins (e.g. PF16 and FAP216 from the C1a-e-c supercomplex, and FAP46 and FAP74 from the C1d projection) that cause severely impaired motility or paralyzed cilia (Brown et al., 2012; Dutcher et al., 1984; Fu et al., 2019; Smith and Lefebvre, 1996). This likely means that the C1b projection, including the FAP246-FAP413-FAP42 subcomplex, is not essential for transmission of the signal that regulates dynein activity and thus ciliary beating. However, *fap246* cells, but not *fap42* cells, frequently changed swimming direction, resulting in curving and even spiraling swimming paths (Fig. 1E). This is typically observed when the two *Chlamydomonas* cilia beat asynchronously, suggesting that FAP246 may play some role in coordinating and/or timing the oscillatory switching of dynein activity to maintain synchrony between the two beating cilia (Polin et al., 2009).

FAP413 has no close homologs in holozoans (Zhao et al., 2020). However, secondary structure prediction suggests that it has a large WD40-repeat domain at the C-terminus with at least 11 WD40 repeats (Fig. S1E). WD40-repeat proteins are a large family found in all eukaryotes and are implicated in a variety of functions ranging from signal transduction and transcription regulation to cell-cycle control and apoptosis. WD40-repeat motifs are thought to act as a site for protein-protein interaction, and proteins containing WD40 repeats are known to serve as platforms for the assembly of protein complexes (Jain and Pandey, 2018). Mutations of WD40-repeat proteins are associated with various human diseases including ciliopathies (Kim and Kim, 2020). Our previous proteomic study predicted that FAP413 is associated with the C1 microtubule but could not unambiguously assign it to one of the projections, because it was significantly decreased in both *fap246* and *fap76* axonemes (Zhao et al., 2019). FAP76 is a C1a protein and part of the C1a-e-c supercomplex (Fu et al., 2019). Therefore, it is possible that another copy of FAP413 is present in the C1a-e-c supercomplex, serving as a binding platform like many WD40-repeat proteins that are known to be promiscuous interactors (Stirnimann et al., 2010).

### The FAP246-FAP413-FAP42 subcomplex is part of a large interconnected CA network

A recurring feature of CA projections and subunits is their high connectivity within the projection and with neighboring CA structures, resulting in a massive CA protein network. For example, we previously reported that the C1a-e-c supercomplex forms multiple connections within the complex and at least three connections with the neighboring C1d projection (Fu et al., 2019), and here we showed that the C1b projection has four internal connections and six connections with neighboring structures, including with C1f and the C1-C2 bridge (Fig. 5). 3D- classification analyses revealed that loss of any protein of the FAP246-FAP413-FAP42 subcomplex caused positional flexibility and/or partial reduction of the neighboring C1b pillar structure and C1f projection. The latter is not in direct contact with the beam-bracket (FAP246-FAP413-FAP42) subcomplex, and thus is likely affected through its connections with the C1b pillar, suggesting a C1b-f supercomplex. Taken together with previous findings that the C1f projection also interacts with the C1d projection, because mutation of the C1d protein FAP74 in *fap7*4 led to loss of both the C1d projection and the C1f projection (previously termed “sheath” in conventional transmission electron microscope studies) (Brown et al., 2012), and that the C1d projection interacts with the C1c protein FAP76 from the C1a-e-c supercomplex (Fu et al., 2019), a picture of a highly interconnected C1a-e-c-d-f-b network emerges.

In our cryo-ET averages of the WT CA, the tip of the C1b beam (FAP42) appears to also connect with corresponding structures of the C2b projection. Previous studies also suggested that the C1b and C2b projections physically interact; specifically, isolated *cpc1* axonemes lacking the C1b projection also frequently lacked the C2b projection (Mitchell and Sale, 1999), and knockdown of hydin, a C2b protein, caused loss of the C2b projection and, in some instances, destabilization of the C1b projection (Lechtreck and Witman, 2007). However, our MS/MS analyses of *fap42* axonemes did not show a decrease of known C2b proteins (Table S1), and a 3D-classification analysis of the C2b projection in *fap42* did not show obvious positional instability or reduction of the projection despite loss of its interface with the C1b beam structure.

### The FAP246-FAP413-FAP42 subcomplex provides mechanical support

In straight flagella, the nine DMTs are cylindrically arranged, and the RSs, which project towards the center of the axoneme, end with their heads on a virtual cylinder with ∼40 nm radius. The two parallel CA microtubules themselves could not interact with all radial spokes at the same time, but the different CA projections vary considerably in length, angle and shape, forming a dense cylindrical CA protein network with ∼40 nm radius that could interact with the RSs from all nine DMTs (Nicastro et al., 2005). At the current cryo-ET resolution, there appears to be an about 5 nm gap between the periphery of the CA projections and the RS heads in straight (inactive) axonemes (Oda et al., 2014). In both forward and backwards swimming *Chlamydomonas*, the diameter of the axoneme is greater in the plane of bend of bent regions than in straight regions of the flagellum, presumably due to transverse stress acting across the axoneme during bending (Fig. S3) (Lindemann and Mitchell, 2007). In the bent regions, the two central microtubules, which in *Chlamydomonas* rotate within the ring of 9 DMTs during flagellar beating, are both in the plane of bend, with the C1 microtubule (predominantly) facing DMT 1, at least in principal bends (Mitchell, 2003). The lateral compression of the axoneme in bent regions implies that the RSs mechanically push against the CA projections in a direction perpendicular to the bend plane; i.e. the C1a-C2a and C1b-C2b projections would be subject to the greatest compression force by the RSs in the bent region (Fig. S3).

Unlike other projections, such as C1c and C1d that are short and closely associated with the C1 microtubule, the C1b beam, which would be closest to the RS heads, is located at the periphery of C1b, 20 nm from the C1-C2 bridge, and is not directly supported by the C1 microtubule. However, the structure of C1b appears to be ideally suited to resist compressive force. The shape of the projection resembles that of a pillar-beam construction, which in architecture is used to transmit load imposed on a horizontal beam (e.g. due to the weight of the roof) to a vertical pillar and floor, which are equipped to resist the compression force. In the case of the C1b projection, the compression force imposed by the RSs onto the FAP42 beam could be transmitted through the connection with the pillar and the FAP246-FAP413 bracket onto the pillar and ultimately the C1 microtubule – both of which should provide more resistance to inwardly directed compressive force. In addition, neighboring beams are connected to each other at the beam tip (close to where the beam interacts with C2b) (Fig. 5A and B; Movie S4), which could distribute locally imposed force by RSs across the sheet-like beam network. Defects in the FAP246-FAP413-FAP42 subcomplex resulted in destabilization of the remaining C1b structures and the C1f projection even in isolated inactive axonemes, i.e. without compression force imposed by bending, suggesting that the FAP246-FAP413-FAP42 subcomplex does indeed play a role in providing mechanical strength and support. The motility defect in the *fap42* and *fap246* mutants may, in part, reflect a transient abnormal compression-induced partial collapse of bent regions of the axoneme during each beat cycle.

Defects in C1b also may affect the regulation of dynein arm activity. Previous studies have shown that mechanical interactions between the CA and RS heads are critical for ciliary motility. For example, Oda and colleagues showed that in the paralyzed *Chlamydomonas* mutant *pf6*, which lacks most of the C1a-e projections, extending the RS heads by addition of a non-ciliary (non-specific) protein of suitable size (e.g. a BCCP-tag) was sufficient to restore flagellar motility (Oda et al., 2014). Oda and colleagues proposed that the C1a projection interacted mechanically and non-specifically with the RSs of DMTs 2, 3 and 4 to transmit signals to the dynein arms. In the *fap42* and *fap246* mutants studied here, loss of the C1b bracket and beam could allow the entire CA to move away from the RSs of DMTs 2, 3, and 4 in bent regions, partially impairing any CA-RS mechano-signaling required to activate or inactivate the dynein arms in those regions. The positions of specific projections relative to the DMTs undergoing active sliding in bent regions could explain why loss of different CA projections has different effects on flagellar motility (Brown et al., 2012; DiPetrillo and Smith, 2011; Lechtreck and Witman, 2007; Mitchell and Sale, 1999; Rupp et al., 2001; Yokoyama et al., 2004; Zhao et al., 2019). Thus the C1b projection might not only support structural integrity and stability of the CA and axoneme, but the mechanical resistance that it provides might be necessary for proper mechano-signaling between the CA and RSs to regulate dynein activity and thus ciliary beating.

## Materials and Methods

### Bioinformatics

Domain predictions of FAP246, FAP42 and FAP413 were performed using PROSITE webserver (http://prosite.expasy.org/) (Sigrist et al., 2013). Sequence alignments were performed using BLAST tool from NCBI (Altschul et al., 1990).

### Strains and culture conditions

*Chlamydomonas reinhardtii* WT strains used were g1 (*nit1*, *agg1*, *mt+*; CC-5415, *Chlamydomonas* Resource Center, https://www.chlamycollection.org) and A54-e18 (CC-2929). A *fap246* mutant strain was originally obtained from the CLiP collection (CLiP ID: LMJ.RY0402.135524) via the *Chlamydomonas* Resource Center and then crossed to g1; a progeny (135524-8A) was named *fap246-1* and used here as in our previous study (Zhao et al., 2019). Another CLiP mutant (LMJ.RY0402.2059300) with an insertion in exon 7 of the *fap42* gene (Li et al., 2016) also was obtained from the *Chlamydomonas* Resource Center; analysis by PCR (Fig. S1) revealed that the cells originally supplied were genetically heterogeneous, so they were cloned, and a clone harboring the insertion was named *fap42-1* and used in subsequent experiments. As previously described (Fu et al., 2018), *Chlamydomonas* cells were maintained in solid Tris-acetate-phosphate (TAP) plates (supplied with 7.5 µg/ml paromomycin for CLiP mutants) and cultured in liquid TAP medium or modified M medium (Witman, 1986) under a 12:12-h light:dark cycle at 23°C with filtered air bubbling into the growth culture. Insertion sites of CLiP mutant *fap42* were confirmed by PCR using the primers listed in Table S4 (Fig. S1F).

### Axoneme preparation

Axonemes of *Chlamydomonas* cells were purified by the pH-shock method as previously described (Song et al., 2015; Witman, 1986). Briefly, cells were cultured in liquid TAP medium, harvested by centrifugation (2,200 rpm for 5 min), and washed twice with fresh M-N/5 minimal medium (Iomini et al., 2009). The cell pellet was resuspended in pH shock buffer (10 mM HEPES, 1 mM SrCl_2_, 4% sucrose, and 1 mM DTT, pH 7.4), and 0.5 M acetic acid was added to the buffer to adjust the pH to 4.3. After 80 s, 1 M KOH was added to increase the pH to 7.2; the pH shock treatment was performed on ice. After pH shock, 5 mM MgSO_4_, 1 mM EGTA, 0.1 mM EDTA, and 100 µl protease inhibitor PMSF (Sigma-Aldrich) were added to the solution. The solution was centrifuged (1800 g for 10 min, 4°C) to separate the detached flagella from the cell bodies. To further purify the flagella, the flagella-containing supernatant was centrifuged twice with a 20% sucrose cushion (2,400 g for 10 min, 4°C). After centrifugation, 1% IGEPAL CA-630 (Sigma-Aldrich) was added to the supernatant for 20 min at 4°C to demembranate the flagella. Axonemes were collected by centrifugation (10,000 g for 10 min, 4°C), and the freshly isolated axonemes were resuspended in HMEEK buffer (30 mM Hepes, 25 mM KCl, 5 mM MgSO_4_, 0.1 mM EDTA, and 1 mM EGTA, pH 7.2). Axonemal samples were either plunge-frozen for cryo-ET or stored at −80°C for biochemical assays or MS.

### Analysis of motility phenotypes

All observations and recordings were performed on cells grown in TAP medium at room temperature. To analyze swimming speed, 100 µl of *Chlamydomonas* cell culture were transferred to a plastic chamber (0.127-mm-deep Fisherbrand UriSystem DeciSlide; Thermo Fisher Scientific). Cells were imaged with a Nikon ECLIPSE LV-N microscope equipped with an Andor Zyla sCMOS Camera. The objective lens used was a Nikon CFI S Plan Fluor ELWD 20x NA 0.45 with a working distance of 8.2-6.9 mm. Movies were recorded at 100 frame/s using NIS-Elements AR (Nikon) software. Swimming speeds were determined using the measurement tools in NIS-Elements AR software. The TrackMate plugin (Tinevez et al., 2017) in Fiji ImageJ was used to trace the swimming trajectory of the cells.

### Liquid chromatography–MS/MS

Axonemal proteins (40 µg) of the WT, *fap42* and *fap246* strains were separated on a 4–12% gradient SDS-polyacrylamide gel (Genscript Biotech). After the dye front migrated on the gel for 3.0–3.5 cm, the gel was stained with Coomassie brilliant blue for 30 min and destained until the background was clear. Each gel lane was cut into four slices, and each slice was further excised into 1-mm cubic pieces. In-gel trypsin digestion, LC-MS/MS and peptide identification were conducted by the proteomics core facility at the University of Texas Southwestern Medical Center. Briefly, protein gel pieces were digested overnight with trypsin (Pierce) following reduction and alkylation with DTT and iodoacetamide (Sigma–Aldrich). The samples then underwent solid-phase extraction cleanup with a Oasis HLB μElution plate (Waters) and the resulting samples were analyzed by LC/MS/MS, using an Orbitrap Fusion Lumos mass spectrometer (Thermo Electron) coupled to an Ultimate 3000 RSLC-Nano liquid chromatography system (Dionex). Samples were injected onto a 75-µm i.d., 75-cm long EasySpray column (Thermo), and eluted with a gradient from 0-28% buffer B over 90 min. Buffer A contained 2% (v/v) acetonitrile and 0.1% formic acid in water, and buffer B contained 80% (v/v) acetonitrile, 10% (v/v) trifluoroethanol, and 0.1% formic acid in water. The mass spectrometer operated in positive ion mode with a source voltage of 1.8 kV and an ion transfer tube temperature of 275 °C. MS scans were acquired at 120,000 resolution in the Orbitrap and up to 10 MS/MS spectra were obtained in the ion trap for each full spectrum acquired using higher-energy collisional dissociation (HCD) for ions with charges 2-7. Dynamic exclusion was set for 25 s after an ion was selected for fragmentation. Raw MS data files were converted to a peak list format and analyzed using the central proteomics facilities pipeline (CPFP), version 2.0.3 (Trudgian and Mirzaei, 2012; Trudgian et al., 2010). Peptide identification was performed using the X!Tandem (Craig and Beavis, 2004) and Open MS Search Algorithm (OMSSA) (Geer et al., 2004) search engines against the *Chlamydomonas reinhardtii* protein database, with common contaminants and reversed decoy sequences appended (Elias and Gygi, 2007). Fragment and precursor tolerances of 10 ppm and 0.5 Da were specified, and three missed cleavages were allowed. Carbamidomethylation of Cys was set as a fixed modification and oxidation of Met was set as a variable modification. Label-free quantitation of proteins across samples was performed using SINQ normalized spectral index software (Trudgian et al., 2011).

### Cryo-ET

Freshly prepared axonemal samples (30 µl) were gently mixed with 10 µl of 10-fold-concentrated, BSA-coated 10-nm gold solution (Sigma Aldrich). 4 µl of the solution was applied to a glow-discharged (30s at 35 mA) copper R2/2 holey carbon grid (Quantifoil Micro Tools). After removing excess liquid by blotting the grid from the back side with Whatman filter paper for 2 s, the grid was plunge-frozen into liquid ethane using a homemade plunge-freezer. Grids were then stored in liquid nitrogen until use.

Grids were loaded into a Titan Krios transmission electron microscope (Thermo Fisher Scientific) operated at 300 kV. The microscope control software SerialEM (Mastronarde, 2005) was used to acquire tilt series images in low-dose mode from −60° to 60° with 2° increments using a dose symmetric tilting scheme (Hagen et al., 2017). For WT axonemes, the images were recorded with a 4,000 × 4,000 K2 DDD camera (Gatan) in counting mode (15 frames, 0.4-s exposure time per frame, dose rate of 8 electrons/pixel/s for each tilt image). For *fap246* and *fap42* axonemes, the images were recorded with a 5,760 x 4,092 K3 DDD camera (Gatan) in counting mode (10 frames, 0.05-s exposure time per frame, dose rate of 26 electrons/pixel/s for each tilt image). For all strains, the post-column energy filter (Gatan) was operated in zero-loss mode (20-eV slit width), and a VPP (Danev et al., 2014) was used with −0.5-µm defocus. The magnification was set to 26,000 with an effective pixel size of 3.2 Å (K3 camera) or 5.5 Å (K2 camera). The total electron dose per tilt series was limited to ∼100 e/ Å^2^.

### Image processing

In brief, motion correction of the frames was done with MotionCorr2 (Zheng et al., 2017). IMOD software (Kremer et al., 1996) was used to align the tilt serial images using the 10-nm gold particles as fiducial markers and to reconstruct the tomograms by the weighted back-projection (WBP) approach. For subtomogram averaging, either CA or DMT repeats were picked from the raw tomograms, and alignment and missing-wedge compensated averaging were performed using PEET software (Nicastro et al., 2006), which is integrated in the IMOD software package. The axoneme and CA orientation (proximal to distal) was determined for each tomogram based on both DMT orientation in the axoneme cross section and initial averages that included only the CA repeats within each tomogram. After the CA polarity and the same center of the repeat were determined for each tomogram, a second alignment was performed combining the CA repeats from all tomograms in the correct orientation and periodicity register. After global alignment of all CA repeats, the alignment of each individual CA microtubule was refined by local alignment of each microtubule and its associated projections separately, while the other microtubule was masked as described previously (Carbajal-Gonzalez et al., 2013; Fu et al., 2019). Visualization of the 3D structures of the averaged CA repeats was done with the UCSF Chimera package software (Pettersen et al., 2004). For generating isosurface renderings, the same isosurface threshold was applied to the WT and mutant averages. Mass estimations of the protein complexes and subvolumes were calculated using the average density of 1.43 g/cm^3^ for proteins (Quillin and Matthews, 2000) and after normalizing the isosurface-rendering threshold to the mass of microtubules in Chimera. Classification analyses used a principal component analysis (PCA) clustering method incorporated in the PEET software (Heumann et al., 2011). The number of tomograms and CA repeats, as well as the estimated resolution of the averages (using FSC 0.5 criterion), are summarized for each strain in Table S5.

## Supplementary materials

Supplementary materials contain three figures, five tables and four movies. Figure S1 shows the protein domain predictions and sequence alignments of FAP246, FAP42 and FAP413; the genotype analysis of the *Chlamydomonas fap42-1* mutant strain; and resolution estimation of the averaged CA repeats of WT, *fap246* and *fap42* axonemes. Figure S2 shows full isosurface renderings of the averaged CA repeats of WT, *fap246* and *fap42* axonemes. Figure S3 shows the proposed model of C1b-RS mechano-signaling that regulates the dynein activities and cilia bending. Table S1 lists the MS/MS results for the axonemes of *fap246* and *fap42* mutants focused on proteins predicted to be novel CA components by two recent proteomic studies (Dai et al., 2020; Zhao et al., 2019) but not yet localized to a specific CA projection. Table S2 lists the exclusive unique FAP246 and FAP413 peptides identified in the proteomic analyses of *fap246* axonemes. Table S3 lists the exclusive unique FAP42 and FAP413 peptides identified in the proteomic analyses of *fap42* axonemes. Table S4 lists the primers used in this study to confirm the mutant insertion site. Table S5 is the summary of image processing information for the strains used in this study. Movie S1 shows the animated 3D visualization of the averaged repeats of the WT *Chlamydomonas* CA. Movie S2 (related to Fig. 2) shows the tomographic slices of the averaged CA repeats of 4 class averages of *fap246* viewed in the longitudinal orientation. Movie S3 (related to Fig. 4) shows the tomographic slices of the averaged CA repeats of 3 class averages of *fap42* viewed in the longitudinal orientation. Movie S4 is the summary animation showing the precise locations of FAP246, FAP413, and FAP42 within the C1b subcomplex.

## Acknowledgments

We thank Dr. Andrew Lemoff and the proteomics core facility at the University of Texas Southwestern Medical Center (UTSW) for the liquid chromatography-MS/MS analyses. We are grateful to Dr. Daniel Stoddard for management of the UTSW cryo-electron microscope facility, which is funded in part by a Cancer Prevention and Research Institute of Texas Core Facility Award (RP170644). This research was supported in part by the computational resources provided by the BioHPC supercomputing facility located in the Lyda Hill Department of Bioinformatics, UT Southwestern Medical Center.

## Data availability

The averaged 3D structures of the CA focused on the C1b projection from different strains have been deposited in the Electron Microscopy Data Bank under accession codes EMD-22648 (WT), EMD-22649 (*fap246*), and EMD-22651 (*fap42*).

## References

Afzelius, B. A. (2004). Cilia-related diseases. J Pathol 204, 470–7.

Altschul, S. F., Gish, W., Miller, W., Myers, E. W. and Lipman, D. J. (1990). Basic local alignment search tool. J Mol Biol 215, 403–10.

Braun, D. A. and Hildebrandt, F. (2017). Ciliopathies. Cold Spring Harb Perspect Biol 9.

Brown, J. M., Dipetrillo, C. G., Smith, E. F. and Witman, G. B. (2012). A FAP46 mutant provides new insights into the function and assembly of the C1d complex of the ciliary central apparatus. J Cell Sci 125, 3904–13.

Brown, J. M. and Witman, G. B. (2014). Cilia and Diseases. Bioscience 64, 1126–1137.

Carbajal-Gonzalez, B. I., Heuser, T., Fu, X., Lin, J., Smith, B. W., Mitchell, D. R. and Nicastro, D. (2013). Conserved structural motifs in the central pair complex of eukaryotic flagella. Cytoskeleton (Hoboken) 70, 101–120.

Craig, R. and Beavis, R. C. (2004). TANDEM: matching proteins with tandem mass spectra. Bioinformatics 20, 1466–7.

Dai, D., Ichikawa, M., Peri, K., Rebinsky, R. and Bui, K. (2019). Identification and mapping of central pair proteins by proteomic analysis. bioRxiv, 739383.

Dai, D., Ichikawa, M., Peri, K., Rebinsky, R. and Bui, K. (2020). Identification and mapping of central pair proteins by proteomic analysis. Biophysics and Physicobiology 17 71–85.

Danev, R., Buijsse, B., Khoshouei, M., Plitzko, J. M. and Baumeister, W. (2014). Volta potential phase plate for in-focus phase contrast transmission electron microscopy. Proc Natl Acad Sci U S A 111, 15635–40.

DiPetrillo, C. G. and Smith, E. F. (2010). Pcdp1 is a central apparatus protein that binds Ca(2+)-calmodulin and regulates ciliary motility. J Cell Biol 189, 601–12.

DiPetrillo, C. G. and Smith, E. F. (2011). The Pcdp1 complex coordinates the activity of dynein isoforms to produce wild-type ciliary motility. Mol Biol Cell 22, 4527–38.

Dutcher, S. K., Huang, B. and Luck, D. J. (1984). Genetic dissection of the central pair microtubules of the flagella of Chlamydomonas reinhardtii. J Cell Biol 98, 229–36.

Dymek, E. E. and Smith, E. F. (2007). A conserved CaM- and radial spoke associated complex mediates regulation of flagellar dynein activity. J Cell Biol 179, 515–26.

Elias, J. E. and Gygi, S. P. (2007). Target-decoy search strategy for increased confidence in large-scale protein identifications by mass spectrometry. Nat Methods 4, 207–14.

Fu, G., Wang, Q., Phan, N., Urbanska, P., Joachimiak, E., Lin, J., Wloga, D. and Nicastro, D. (2018). The I1 dynein-associated tether and tether head complex is a conserved regulator of ciliary motility. Mol Biol Cell 29, 1048–1059.

Fu, G., Zhao, L., Dymek, E., Hou, Y., Song, K., Phan, N., Shang, Z., Smith, E. F., Witman, G. B. and Nicastro, D. (2019). Structural organization of the C1a-e-c supercomplex within the ciliary central apparatus. J Cell Biol 218, 4236–4251.

Geer, L. Y., Markey, S. P., Kowalak, J. A., Wagner, L., Xu, M., Maynard, D. M., Yang, X., Shi, W. and Bryant, S. H. (2004). Open mass spectrometry search algorithm. J Proteome Res 3, 958–64.

Gui, L., Song, K., Tritschler, D., Bower, R., Yan, S., Dai, A., Augspurger, K., Sakizadeh, J., Grzemska, M., Ni, T. et al. (2019). Scaffold subunits support associated subunit assembly in the Chlamydomonas ciliary nexin-dynein regulatory complex. Proc Natl Acad Sci U S A.

Hagen, W. J. H., Wan, W. and Briggs, J. A. G. (2017). Implementation of a cryo-electron tomography tilt-scheme optimized for high resolution subtomogram averaging. J Struct Biol 197, 191–198.

Heumann, J. M., Hoenger, A. and Mastronarde, D. N. (2011). Clustering and variance maps for cryo-electron tomography using wedge-masked differences. J Struct Biol 175, 288–299.

Iomini, C., Till, J. E. and Dutcher, S. K. (2009). Genetic and phenotypic analysis of flagellar assembly mutants in Chlamydomonas reinhardtii. Methods Cell Biol 93, 121–43.

Jain, B. P. and Pandey, S. (2018). WD40 Repeat Proteins: Signalling Scaffold with Diverse Functions. Protein J 37, 391–406.

Khan, N., Shah, P. P., Ban, D., Trigo-Mourino, P., Carneiro, M. G., DeLeeuw, L., Dean, W. L., Trent, J. O., Beverly, L. J., Konrad, M. et al. (2019). Solution structure and functional investigation of human guanylate kinase reveals allosteric networking and a crucial role for the enzyme in cancer. J Biol Chem 294, 11920–11933.

Kikkawa, M. (2013). Big steps toward understanding dynein. J Cell Biol 202, 15–23.

Kim, Y. and Kim, S. H. (2020). WD40-repeat proteins in ciliopathies and congenital disorders of endocrine system. Endocrinol Metab (Seoul) 35, 494–506.

Kremer, J. R., Mastronarde, D. N. and McIntosh, J. R. (1996). Computer visualization of three-dimensional image data using IMOD. J Struct Biol 116, 71–6.

Lechtreck, K. F. and Witman, G. B. (2007). Chlamydomonas reinhardtii hydin is a central pair protein required for flagellar motility. Journal of Cell Biology 176, 473–482.

Lee, L., Campagna, D. R., Pinkus, J. L., Mulhern, H., Wyatt, T. A., Sisson, J. H., Pavlik, J. A., Pinkus, G. S. and Fleming, M. D. (2008). Primary ciliary dyskinesia in mice lacking the novel ciliary protein Pcdp1. Molecular and Cellular Biology 28, 949–57.

Li, X., Zhang, R., Patena, W., Gang, S. S., Blum, S. R., Ivanova, N., Yue, R., Robertson, J. M., Lefebvre, P. A., Fitz-Gibbon, S. T. et al. (2016). An Indexed, Mapped Mutant Library Enables Reverse Genetics Studies of Biological Processes in Chlamydomonas reinhardtii. Plant Cell 28, 367–87.

Lin, J. and Nicastro, D. (2018). Asymmetric distribution and spatial switching of dynein activity generates ciliary motility. Science 360.

Lindemann, C. B. and Mitchell, D. R. (2007). Evidence for axonemal distortion during the flagellar beat of Chlamydomonas. Cell Motil Cytoskeleton 64, 580–9.

Liu, Y., DeBoer, K., de Kretser, D. M., O’Donnell, L., O’Connor, A. E., Merriner, D. J., Okuda, H., Whittle, B., Jans, D. A., Efthymiadis, A. et al. (2015). LRGUK-1 is required for basal body and manchette function during spermatogenesis and male fertility. Plos Genetics 11, e1005090.

Loreng, T. D. and Smith, E. F. (2017). The Central Apparatus of Cilia and Eukaryotic Flagella. Cold Spring Harb Perspect Biol 9.

Mastronarde, D. N. (2005). Automated electron microscope tomography using robust prediction of specimen movements. J Struct Biol 152, 36–51.

McKenzie, C. W., Craige, B., Kroeger, T. V., Finn, R., Wyatt, T. A., Sisson, J. H., Pavlik, J. A., Strittmatter, L., Hendricks, G. M., Witman, G. B. et al. (2015). CFAP54 is required for proper ciliary motility and assembly of the central pair apparatus in mice. Mol Biol Cell 26, 3140–9.

Mitchell, B. F., Pedersen, L. B., Feely, M., Rosenbaum, J. L. and Mitchell, D. R. (2005). ATP production in Chlamydomonas reinhardtii flagella by glycolytic enzymes. Mol Biol Cell 16, 4509–18.

Mitchell, D. R. and Sale, W. S. (1999). Characterization of a Chlamydomonas insertional mutant that disrupts flagellar central pair microtubule-associated structures. J Cell Biol 144, 293–304.

Nakazawa, Y., Ariyoshi, T., Noga, A., Kamiya, R. and Hirono, M. (2014). Space-Dependent Formation of Central Pair Microtubules and Their Interactions with Radial Spokes. PLoS One 9.

Nicastro, D., McIntosh, J. R. and Baumeister, W. (2005). 3D structure of eukaryotic flagella in a quiescent state revealed by cryo-electron tomography. Proc Natl Acad Sci U S A 102, 15889–94.

Nicastro, D., Schwartz, C., Pierson, J., Gaudette, R., Porter, M. E. and McIntosh, J. R. (2006). The molecular architecture of axonemes revealed by cryoelectron tomography. Science 313, 944–8.

Oda, T., Yanagisawa, H., Yagi, T. and Kikkawa, M. (2014). Mechanosignaling between central apparatus and radial spokes controls axonemal dynein activity. J Cell Biol 204, 807–19.

Olbrich, H., Schmidts, M., Werner, C., Onoufriadis, A., Loges, N. T., Raidt, J., Banki, N. F., Shoemark, A., Burgoyne, T., Al Turki, S. et al. (2012). Recessive HYDIN mutations cause primary ciliary dyskinesia without randomization of left-right body asymmetry. American Journal of Human Genetics 91, 672–84.

Pettersen, E. F., Goddard, T. D., Huang, C. C., Couch, G. S., Greenblatt, D. M., Meng, E. and Ferrin, T. E. (2004). UCSF Chimera--a visualization system for exploratory research and analysis. J Comput Chem 25, 1605–12.

Polin, M., Tuval, I., Drescher, K., Gollub, J. P. and Goldstein, R. E. (2009). Chlamydomonas swims with two “gears” in a eukaryotic version of run-and-tumble locomotion. Science 325, 487–90.

Poprzeczko, M., Bicka, M., Farahat, H., Bazan, R., Osinka, A., Fabczak, H., Joachimiak, E. and Wloga, D. (2019). Rare Human Diseases: Model Organisms in Deciphering the Molecular Basis of Primary Ciliary Dyskinesia. Cells 8.

Quillin, M. L. and Matthews, B. W. (2000). Accurate calculation of the density of proteins. Acta Crystallogr D Biol Crystallogr 56, 791–4.

Rao, V. G., Sarafdar, R. B., Chowdhury, T. S., Sivadas, P., Yang, P., Dongre, P. M. and D’Souza, J. S. (2016). Myc-binding protein orthologue interacts with AKAP240 in the central pair apparatus of the Chlamydomonas flagella. BMC Cell Biol 17, 24.

Roberts, A. J., Kon, T., Knight, P. J., Sutoh, K. and Burgess, S. A. (2013). Functions and mechanics of dynein motor proteins. Nat Rev Mol Cell Biol 14, 713–26.

Rupp, G., O’Toole, E. and Porter, M. E. (2001). The Chlamydomonas PF6 locus encodes a large alanine/proline-rich polypeptide that is required for assembly of a central pair projection and regulates flagellar motility. Mol Biol Cell 12, 739–51.

Sigrist, C. J., de Castro, E., Cerutti, L., Cuche, B. A., Hulo, N., Bridge, A., Bougueleret, L. and Xenarios, I. (2013). New and continuing developments at PROSITE. Nucleic Acids Res 41, D344–7.

Smith, E. F. and Lefebvre, P. A. (1996). PF16 encodes a protein with armadillo repeats and localizes to a single microtubule of the central apparatus in Chlamydomonas flagella. J Cell Biol 132, 359–70.

Smith, E. F. and Yang, P. (2004). The radial spokes and central apparatus: mechano-chemical transducers that regulate flagellar motility. Cell Motil Cytoskeleton 57, 8–17.

Song, K., Awata, J., Tritschler, D., Bower, R., Witman, G. B., Porter, M. E. and Nicastro, (2015). In situ localization of N and C termini of subunits of the flagellar nexin-dynein regulatory complex (N-DRC) using SNAP tag and cryo-electron tomography. J Biol Chem 290, 5341–53.

Stirnimann, C. U., Petsalaki, E., Russell, R. B. and Muller, C. W. (2010). WD40 proteins propel cellular networks. Trends Biochem Sci 35, 565–74.

Teves, M. E., Nagarkatti-Gude, D. R., Zhang, Z. and Strauss, J. F., 3rd. (2016). Mammalian axoneme central pair complex proteins: Broader roles revealed by gene knockout phenotypes. Cytoskeleton (Hoboken) 73, 3–22.

Tinevez, J. Y., Perry, N., Schindelin, J., Hoopes, G. M., Reynolds, G. D., Laplantine, E., Bednarek, S. Y., Shorte, S. L. and Eliceiri, K. W. (2017). TrackMate: An open and extensible platform for single-particle tracking. Methods 115, 80–90.

Trudgian, D. C. and Mirzaei, H. (2012). Cloud CPFP: a shotgun proteomics data analysis pipeline using cloud and high performance computing. J Proteome Res 11, 6282–90.

Trudgian, D. C., Ridlova, G., Fischer, R., Mackeen, M. M., Ternette, N., Acuto, O., Kessler, B. M. and Thomas, B. (2011). Comparative evaluation of labelifree SINQ normalized spectral index quantitation in the central proteomics facilities pipeline. Proteomics 11, 2790–2797.

Trudgian, D. C., Thomas, B., McGowan, S. J., Kessler, B. M., Salek, M. and Acuto, O. (2010). CPFP: a central proteomics facilities pipeline. Bioinformatics 26, 1131–2.

Viswanadha, R., Sale, W. S. and Porter, M. E. (2017). Ciliary Motility: Regulation of Axonemal Dynein Motors. Cold Spring Harb Perspect Biol 9.

Wirschell, M., Nicastro, D., Porter, M.E., and Sale, W.S. (2009). The regulation of axonemal bending. The Chlamydomonas Sourcebook 3, 253–282.

Witman, G. B. (1986). Isolation of Chlamydomonas flagella and flagellar axonemes. Methods Enzymol 134, 280–90.

Witman, G. B., Plummer, J. and Sander, G. (1978). Chlamydomonas flagellar mutants lacking radial spokes and central tubules. Structure, composition, and function of specific axonemal components. J Cell Biol 76, 729–47.

Yokoyama, R., O’Toole, E., Ghosh, S. and Mitchell, D. R. (2004). Regulation of flagellar dynein activity by a central pair kinesin. Proc Natl Acad Sci U S A 101, 17398–403.

Zhang, H. and Mitchell, D. R. (2004). Cpc1, a Chlamydomonas central pair protein with an adenylate kinase domain. J Cell Sci 117, 4179–88.

Zhang, Z., Zariwala, M. A., Mahadevan, M. M., Caballero-Campo, P., Shen, X., Escudier, E., Duriez, B., Bridoux, A. M., Leigh, M., Gerton, G. L. et al. (2007). A heterozygous mutation disrupting the SPAG16 gene results in biochemical instability of central apparatus components of the human sperm axoneme. Biol Reprod 77, 864–71.

Zhao, L., Hou, Y., McNeill, N. A. and Witman, G. B. (2020). The unity and diversity of the ciliary central apparatus. Philos Trans R Soc Lond B Biol Sci 375, 20190164.

Zhao, L., Hou, Y., Picariello, T., Craige, B. and Witman, G. B. (2019). Proteome of the central apparatus of a ciliary axoneme. J Cell Biol 218, 2051–2070.

Zheng, S. Q., Palovcak, E., Armache, J. P., Verba, K. A., Cheng, Y. and Agard, D. A. (2017). MotionCor2: anisotropic correction of beam-induced motion for improved cryo-electron microscopy. Nat Methods 14, 331–332.

